# Bottom-up and top-down computations in high-level visual cortex

**DOI:** 10.1101/053595

**Authors:** Kendrick N. Kay, Jason D. Yeatman

## Abstract

The ability to read a page of text or recognize a person’s face depends on category-selective visual regions in ventral temporal cortex (VTC). To understand how these regions mediate word and face recognition, it is necessary to characterize how stimuli are represented and how this representation is used in the execution of a cognitive task. Here, we show that the response of a category-selective region in VTC can be computed as the degree to which the low-level properties of the stimulus match a category template. Moreover, we show that during execution of a task, the bottom-up representation is scaled by the intraparietal sulcus (IPS), and that the level of IPS engagement reflects the cognitive demands of the task. These results provide a unifying account of neural processing in VTC in the form of a model that addresses both bottom-up and top-down effects and quantitatively predicts VTC responses.

## Main text

How does visual cortex work? One approach to answering this question consists in building functional models that characterize the computations that are implemented by neurons and their circuitry^1,2^. This approach has been fruitful for the front end of the visual system, where relatively simple image computations have been shown to characterize the spiking activity of neurons in the retina, thalamus, and V1^3,4^. Based on this pioneering work in electrophysiology, researchers have extended the modeling approach to characterize responses in human visual cortex, as measured by functional magnetic resonance imaging (fMRI)^5–7^.

Models of early visual processing have been able to offer accurate explanations of low-level perceptual functions such as contrast detection^8,9^ and orientation discrimination^10^. However, these models are insufficient to explain high-level perceptual functions such as the ability to read a page of text or recognize a face. These abilities are believed to depend on category-selective regions in ventral temporal cortex (VTC), but the computations that give rise to category-selective responses are poorly understood.

The goal of the present study is to develop a fully computable model that predicts neural responses in high-level visual cortex of human observers while they perform different cognitive tasks on a wide range of images. Achieving this goal requires four substantial innovations: First, we need to develop a forward model that formally specifies the visual information encoded in the activity of high-level visual regions. Second, we need to dissociate bottom-up stimulus-driven effects from modulation by top-down cognitive processes and characterize how these processes alter the stimulus representation. Third, we need to localize the source of the top-down effects and integrate bottom-up and top-down computations into a single consolidated model. Finally, for completeness, the neural computations should be linked to the measured behavior of the visual observer. In this study, we achieve these four innovations and provide the first comprehensive account of human high-level visual cortex.

## VTC reflects both stimulus properties and cognitive task

Ventral temporal cortex (VTC) is divided into a mosaic of high-level visual regions that respond selectively to specific image categories, and are believed to play an essential role in object perception^11,12^. We focus on two specific VTC regions, the visual word form area (VWFA), which selectively responds to words^13–15^, and the fusiform face area (FFA), which selectively responds to faces^16,17^.

We measured blood oxygenation level dependent (BOLD) responses to a set of carefully controlled images while manipulating the cognitive task that the subjects performed on the stimuli. The first task was designed to minimize the influence of cognitive processes on sensory processing of the stimulus. Subjects performed a demanding perceptual task on a small dot (0.12° × 0.12°) presented at fixation. In this *fixation task*, the presented stimuli are irrelevant to the subject, and we interpret evoked activity as reflecting the intrinsic, bottom-up response from VTC.

To first approximation, much of the variance in bottom-up responses from VWFA and FFA is explained by the category of stimulus (**Figure 1**). However, we find that responses are not invariant to low-level properties of the stimulus: both image contrast and phase coherence modulate response amplitudes. For example, the response to a word in VWFA is 2.4 times stronger when the word is presented at 100% contrast as compared to 3% contrast. The existence and magnitude of these bottom-up effects may be surprising (see also refs. 18,19) as theories of high-level vision are typically predicated on VTC responses being invariant to low-level features^20,21^. Our measurements indicate that an accurate model of the computations performed by VTC must consider not only the stimulus category but also low-level features of the stimulus.

**Figure 1.**
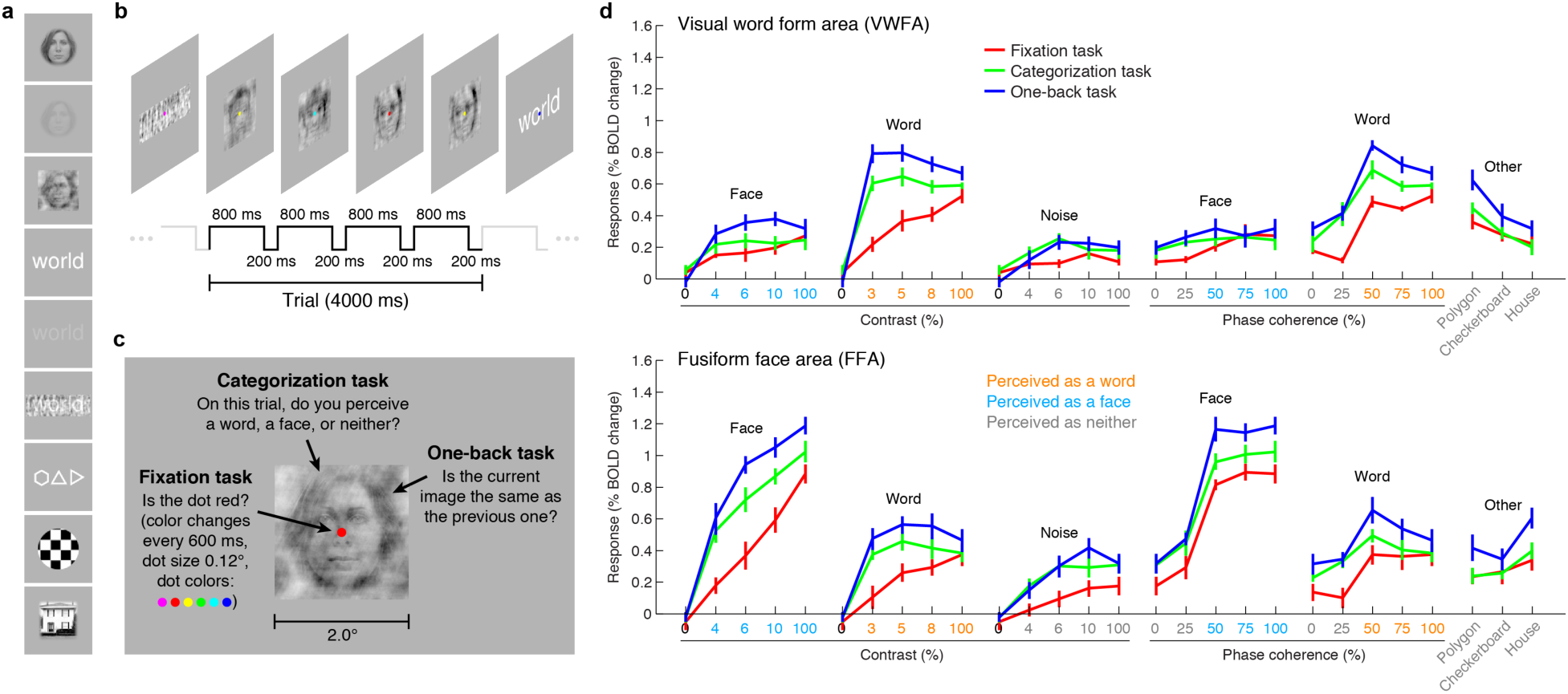
VTC responses depend on both stimulus properties and cognitive task. (a) *Stimuli*. Stimuli included faces, words, and noise patterns presented at different contrasts and phase-coherence levels, as well as full-contrast polygons, checkerboards, and houses. (b) *Trial design*. Each trial consisted of four images drawn from the same stimulus type. (c) *Tasks*. On a given trial, subjects performed one of three tasks. (d) *Evoked responses in VWFA (top) and FFA (bottom) for different stimuli and tasks*. Color of *x*-axis label indicates the perceived stimulus category as reported by the subjects. Error bars indicate bootstrapped 68% CIs.

We also measured VTC responses while subjects performed a *categorization task*, in which the subject reports the perceived category of the stimulus, and a *one-back task*, in which the subject detects consecutive repetitions of stimulus frames. Despite presentation of identical stimuli across the three tasks, there are substantial changes in evoked VTC responses (**Figure 1d**). Responses are larger for the categorization and one-back tasks compared to the fixation task, and we interpret these response increases as reflecting top-down modulation. In some cases, the top-down modulation is even larger than the modulation achieved by manipulation of the stimulus. For example, the VWFA response to 3%-contrast words during the one-back task exceeds the response to 100%-contrast words. Task effects in lower-level areas exist but are smaller in size (**Extended Data Figure 1**). Our measurements indicate that VTC responses cannot be interpreted without specifying the cognitive state of the observer. A complete model of the computations performed by VTC must consider the cognitive task in addition to stimulus properties.

## Top-down modulation acts as a stimulus-specific scaling

Our measurements suggest that high-level visual regions in VTC flexibly adapt to the demands of the cognitive task, but the precise nature of these effects remains unclear. By visualizing VTC responses as points in a multi-dimensional neural space with VWFA, FFA, and hV4 BOLD response amplitudes as the axes, we see that responses to words and faces lie on specific manifolds, appearing as “arms” that emanate from the origin (**Figure 2**). Importantly, we observe that the categorization and one-back tasks act as a scaling mechanism on the representation observed during the fixation task. The scaling mechanism moves the representation of each stimulus along the arms and away from the origin. Moreover, the amount of scaling is not constant across stimuli but is stimulus-specific, and this is most evident when considering the lowest contrast stimuli (**Figure 2**, **black dots**).

**Figure 2.**
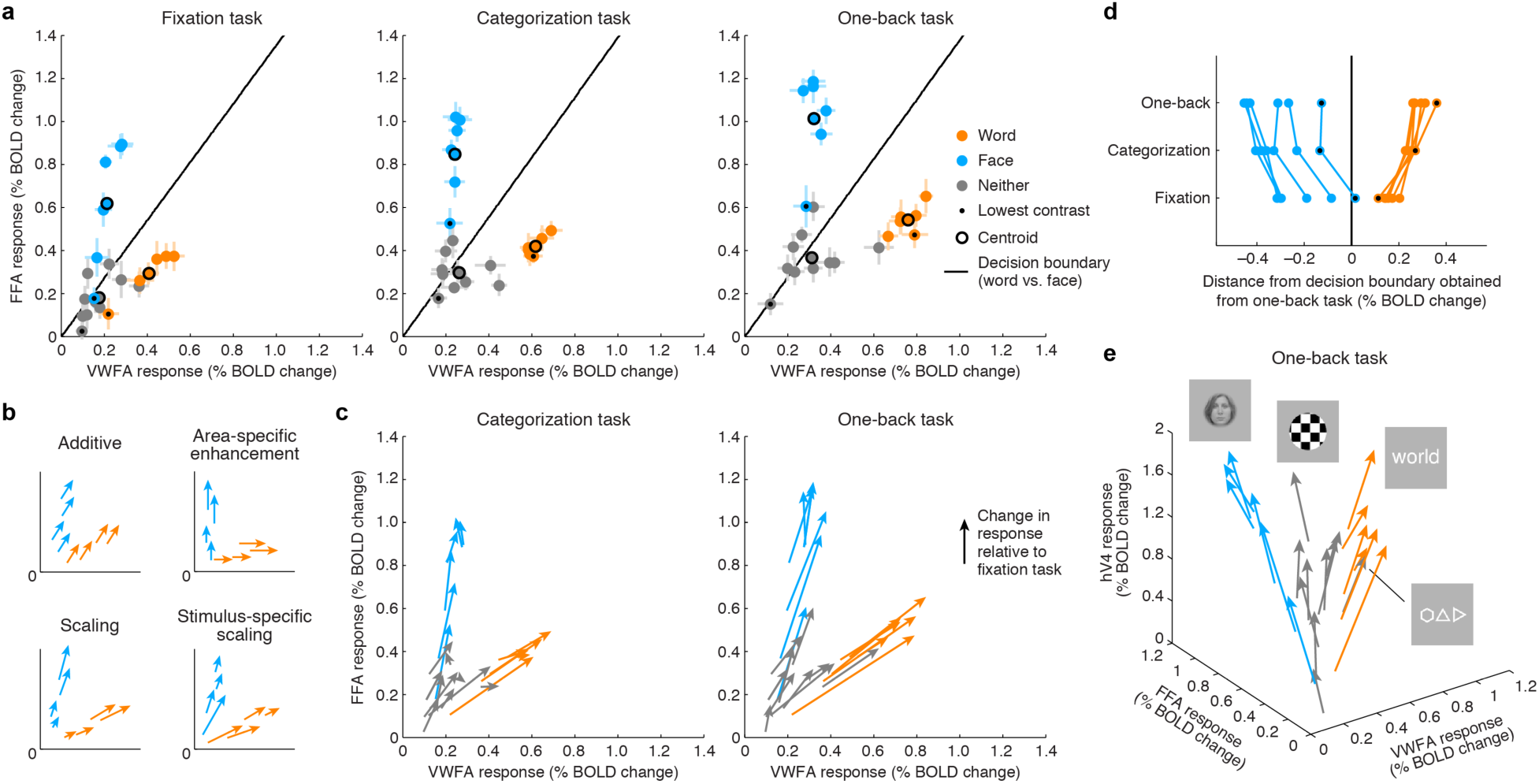
Top-down stimulus-specific scaling of VTC representation. (a) *Responses plotted in multi-dimensional neural space*. Each dot indicates ROI (VWFA, FFA) responses to a stimulus. In each plot, the black line indicates a linear decision boundary separating words and faces (nearest-centroid classifier, angular distance). (b) *Schematics of potential top-down mechanisms*. (c) *Categorization and one-back tasks produce stimulus-specific scaling*. Arrows indicate the change in representation compared to the fixation task. (d) *Scaling improves readout*. Scaling moves neural responses away from the decision boundary, thereby improving signal-to-noise ratio. (e) *Separation of other stimulus categories*. Including hV4 as a third dimension reveals that stimuli categorized as neither words nor faces manifest as a third “arm” that emanates from the origin.

The visualization also shows that substantial responses to non-preferred categories are present in each ROI (e.g., faces in VWFA, words in FFA) and that these responses are scaled during the stimulus-directed tasks. Thus, not only is information regarding non-preferred categories present in each ROI, but this information is actively modulated when subjects perform a perceptual task on those categories. These observations support the view that the brain uses a distributed strategy for perceptual processing and that category-selective regions are components of a more general network of regions that coordinate to extract visual information^22,23^. An alternative scheme, more in line with a modular view of perceptual processing^24,25^, is area-specific enhancement, in which the representation of a stimulus is enhanced only in the region that is selective for that stimulus (e.g., enhancement of words only in VWFA, enhancement of faces only in FFA). This scheme is not supported by our measurements (**Figures 2b** **and** **2c**).

A simple interpretation of the scaling effects is that they serve to increase signal-to-noise ratio in visually evoked responses in VTC^26^. For example, assuming that one use of the stimulus representation in VTC is to discriminate whether the presented stimulus is a word or face (or, more generally, identify the category of the stimulus^27^), the scaling induced by the stimulus-directed tasks serves to increase the distance of neural responses from a linear decision boundary that separates words and faces (**Figure 2c**). Interestingly, the categorization and one-back tasks appear to act via the same scaling mechanism; the stronger scaling observed for the one-back task might be a consequence of either longer-duration or more effortful cognitive processing. These results suggest that cognition has a simple, stereotyped effect on visual cortex: cognitive processes do not impart any additional tuning or selectivity but simply serve to amplify the selectivity that is already computed by visual cortex.

## IPS provides top-down modulation to VTC

To lay the foundation for building a quantitative model that predicts VTC responses, we seek to identify the neural circuitry that generates the observed top-down effects in VTC. There are two candidate mechanisms that might generate the top-down effects. The first is that sensitivity to task is locally generated from the neuronal architecture of VTC itself.We explore an alternative hypothesis whereby top-down modulation is induced by input from another brain region that is sensitive to task demands. To identify this region, we perform a connectivity analysis in which we first subtract the bottom-up signal in VTC, as given by responses measured during the fixation task, from responses measured during the categorization and one-back tasks. We then correlate these residuals, which isolate the top-down signal, against the responses of every cortical location.

Applying this connectivity analysis to our data, we find that responses in the intraparietal sulcus (IPS) predict the top-down enhancement of VTC responses (**Figure 3b**). Hence, our data suggest the IPS is the region that induces task sensitivity in VTC. As a control, if we omit the subtraction step and simply correlate raw VTC responses with the responses of different cortical locations, we find that the correlation is instead strongest with a range of areas spanning occipital cortex (**Figure 3a**). This indicates that the VTC response is a mixture of bottom-up and top-down effects and that the top-down influence from the IPS becomes clear only when bottom-up effects are removed. Comparing our results to a publicly available atlas^28^, we estimate that the source of top-down modulation is localized to the IPS-0 and IPS-1 subdivisions of the IPS (see also **Extended Data Figure 2**).

**Figure 3.**
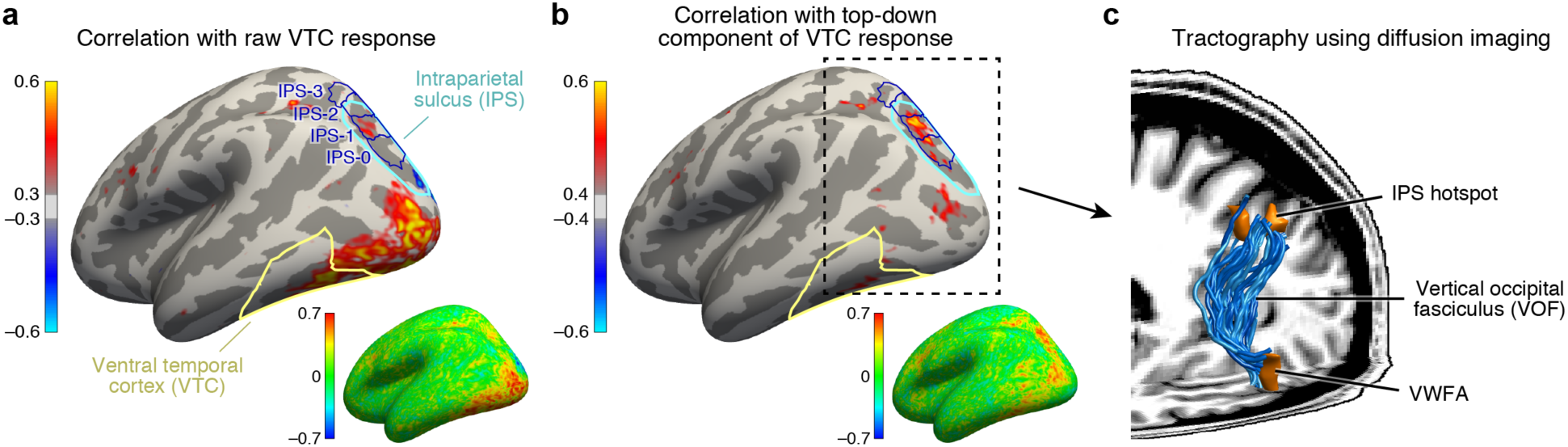
IPS is the source of top-down modulation to VTC. (a) *Correlation with raw VTC response*. This map depicts the correlation between the VTC response observed during the categorization and one-back tasks with the response at each cortical location (inset shows an unsmoothed and unthresholded map). Positive correlations are broadly distributed across occipital cortex. (b) *Correlation with top-down component of VTC response*. After removing bottom-up responses (fixation task), the correlation is spatially localized to a hotspot in IPS-0/1. (c) *Tractography using diffusion MRI*. We find that the vertical occipital fasciculus^31^ connects VWFA and FFA to the IPS hotspot in each subject (rendering shows a representative subject).

Previous research has identified IPS as playing a key role in controlling spatial attention^29,30^. Our results extend these findings by showing that, despite the fact that spatial attention is always directed towards the foveal stimulus during the categorization and one-back tasks, the amount of modulation from the IPS is flexible and varies depending on properties of the stimulus and demands of the task. This mechanism could explain the finding that difficult tasks enhance visual responses^8^. The direct influence of IPS on neural responses in VTC is further supported by the existence of a large white-matter pathway connecting dorsal and ventral visual cortex, called the vertical occipital fasciculus (VOF)^31,32^. In fact, using diffusion-weighted MRI and tractography, we show that the VOF specifically connects the VWFA and FFA with the functionally identified peak region in the IPS for the eight subjects with diffusion data (**Figure 3c**). The VWFA falls within the ventral terminations of the VOF for seven subjects and, for the eighth, the VWFA is 2.7 mm anterior to the VOF, which is well within the margin of error for tractography^33^. The FFA falls within the ventral terminations of the VOF for all eight subjects. These results provide an elegant example of how anatomy subserves function, and sets the stage for a circuit-level computational model that, guided by anatomical constraints, characterizes the computations that emerge from interactions between multiple brain regions.

Guided by our observations of stimulus and task modulations of VTC (**Figure 1**), top-down stimulus-specific scaling (**Figure 2**), and localization of top-down effects to the IPS (**Figure 3**), we now turn to the task of developing a comprehensive model that accounts for our experimental measurements of VTC and IPS. In particular, we seek a general and fully computable model— that is, a model that can operate on any arbitrary visual image and quantitatively predict brain activity and behavior to a high level of accuracy^6,34–36^. Developing such a model is the focus of the rest of this paper.

## Template model of bottom-up VTC response

The first component of our model addresses bottom-up stimulus processing in VTC. Although the field has long understood that stimulus category is a good predictor of evoked responses^16,17^, we do not yet have a computational explanation of this phenomenon. In other words, although we are able to use our own visual systems to assign a label such as “word” or “face” to describe the data, we have not yet identified the computational operations that enable our visual systems to derive these labels in the first place. An additional limitation of our conceptual understanding is that it fails to account for the sensitivity of VWFA and FFA to low-level image properties. We therefore ask: Is it possible to develop a quantitative characterization of the bottom-up computations performed by high-level regions in human VTC?

Extending an existing computational model of fMRI responses in the visual system^6,37,38^, we conceive of a model involving two stages of image computations (**Figure 4**, **upper green box**). The first stage consists of a set of local oriented filters, akin to what has been used to model physiological responses in V1^3,39^. The second stage consists of a normalized dot product applied to the outputs of the first stage. This dot product computes how well a given stimulus matches a category template (e.g., a word template for VWFA, a face template for FFA). This model, termed the *Template model*, is theoretically motivated and is consistent with hierarchical theories of visual processing^1,27,40–42^. Moreover, unlike recently popular deep neural network models that also involve hierarchical processing^35,36,43^, the model we propose is parsimonious with only three free parameters, and is therefore straightforward to fit and interpret (see **Figure 4** and Methods).

**Figure 4.**
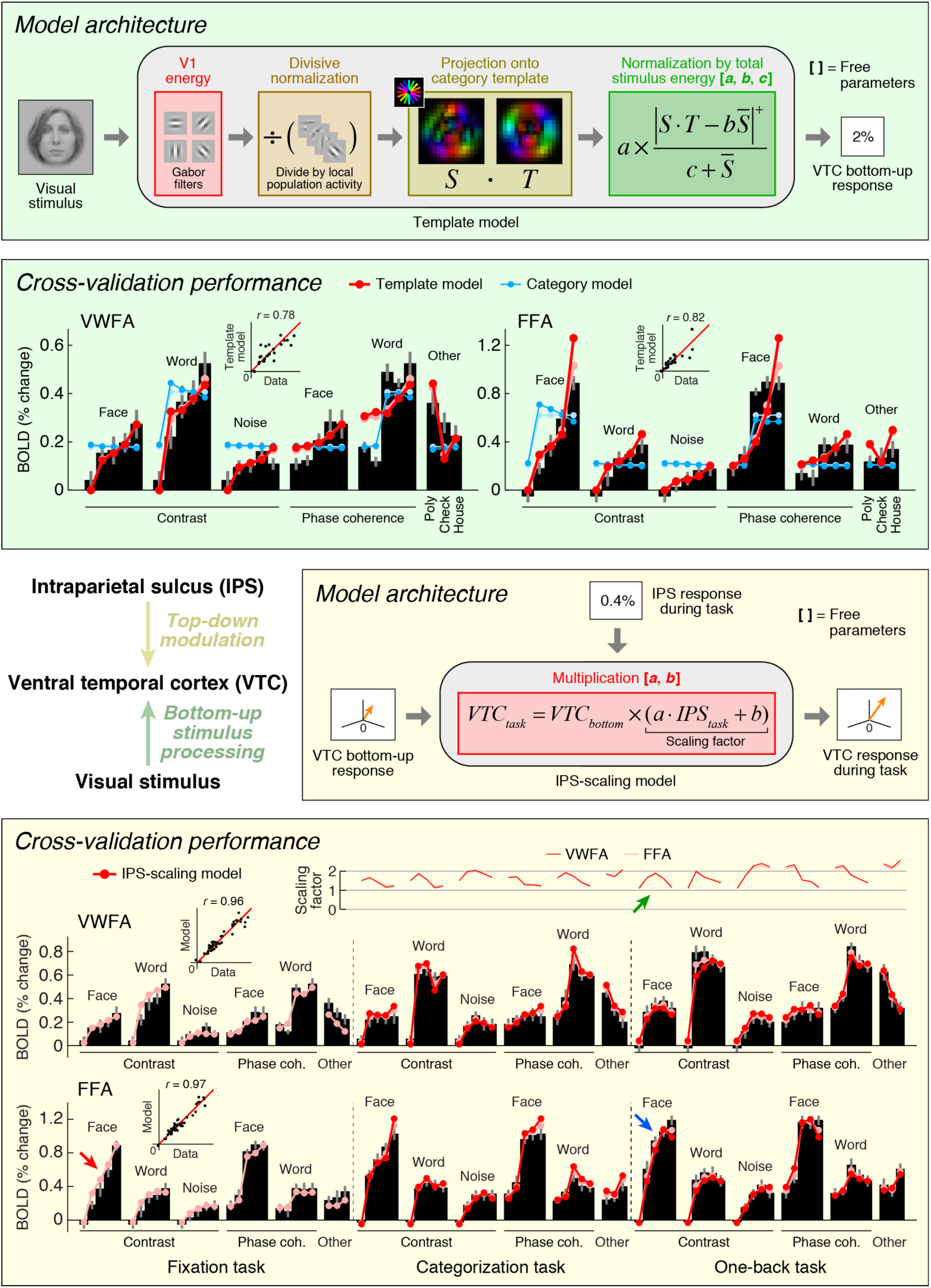
Model of bottom-up and top-down computations in VTC. Bottom-up stimulus-driven responses in VTC (fixation task) can be modeled as a series of image computations (top panels, green). Black bars indicate data, dark lines and dark dots indicate model predictions (leave-one-out cross-validation), and light lines and light dots indicate model fits (no cross-validation). Top-down modulation of VTC responses (categorization task, one-back task) can be modeled as a scaling of the bottom-up responses, with the amount of scaling proportional to the IPS signal (bottom panels, yellow). Scatter plots in the inset compare model predictions against the data. For detailed cross-validation results and control models, see **Extended Data Figure 3**.

Applying the Template model to responses measured during the fixation task, we find that the Template model accurately predicts a large amount of variance in the responses of VWFA and FFA (**Figure 4**, **lower green box**). The model outperforms a phenomenological model, termed the *Category model*, that posits that perceived stimulus category is sufficient to predict the response of category-selective regions. The model also outperforms simpler models that include only one of the two processing stages (**Extended Data Figure 3**). Notably, the Template model is able to predict with substantial accuracy the response to non-preferred stimulus categories in each ROI. This suggests that responses to non-preferred stimuli are meaningful and the result of a well-defined computation performed by the visual system^23^.

The Template model advances us towards a computational understanding of VTC by demonstrating that VTC computes a specific template-matching operation on incoming visual inputs filtered by early visual cortex. Moreover, the model clarifies the nature of high-level visual representations. The present results indicate that although high-level representations are not identical to low-level properties, they are built from, and fundamentally tied to, low-level properties through a series of linear and nonlinear operations. This conclusion is consistent with classic hierarchical theories of visual cortex^1,27,40–42^, and our model can be viewed as a potential mechanism for how semantic tuning properties emerge in visual cortex^34^. Thus, we contend that when studying high-level sensory representations in the brain, a precise characterization of the stimulus still matters.

## IPS-scaling model of VTC modulation

The second component of our model addresses top-down modulation of responses in VTC. Building upon our earlier observations that the top-down modulation is localized to the IPS (see **Figure 3**) and acts as a scaling mechanism on responses in VWFA and FFA (see **Figure 2**), we propose that the magnitude of the IPS response to a stimulus indicates the amount of top-down scaling that is applied to bottom-up sensory responses in VTC (**Figure 4**, **upper yellow box**). We find that this model, termed the *IPS-scaling model*, accurately characterizes the observed data (**Figure 4**, **lower yellow box**). For example, notice that the FFA response to faces increases gradually for each contrast increment during the fixation task (relatively unsaturated contrast-response function, red arrow). When subjects perform the one-back task, we observe a U-shaped contrast-response function in IPS (green arrow); multiplication of the two functions predicts a contrast-response function that is highly saturated and accurately matches the observed contrast-response function in FFA during the one-back task (blue arrow).

Importantly, the IPS-scaling model uses a single set of scale and offset parameters on the IPS response and accurately predicts scaling of VTC responses across the categorization and one-back tasks (**Figure 4**, **lower yellow box, top plot**). This finding suggests that the scaling of VTC by IPS is a general mechanism supporting perception and is independent of the specific cognitive task performed by the observer. Furthermore, the scale and offset parameters that are estimated from the data show that when IPS exhibits close to zero evoked activity (e.g. FACE at 100%-contrast; see **Extended Data Figure 1**), the corresponding scaling factor is close to one. This has a sensible interpretation: when IPS is inactive, we observe only the bottom-up response in VTC and no top-down modulation.

## Drift diffusion model of IPS

Although informative, the finding that IPS provides top-down stimulus-specific scaling of VTC is an incomplete explanation, as the burden of explaining the top-down effects is simply shifted to the IPS. We are thus left wondering: is it possible to explain the response profile of the IPS? In particular, can we explain why the IPS is more active for certain stimuli compared to others? Answering these questions will provide a critical link between cognitive state and IPS activity.

The third and final component of our model offers an explanation of evoked responses in IPS. Inspired by previous research on perceptual decision-making^44–46^, we implement a *Drift diffusion model* that attempts to account for IPS responses measured during the categorization task (**Figure 5**, **upper purple box**). The model uses VTC responses during the fixation task as a measure of sensory evidence, and posits that the IPS accumulates evidence from VTC over time and exhibits an activity level that is monotonically related to accumulation time. For example, when VTC responses are small, as is the case for low-contrast stimuli, sensory evidence for stimulus category is weak, leading to long accumulation times (indexed by measurements of reaction time during the experiment), and large IPS responses.

**Figure 5.**
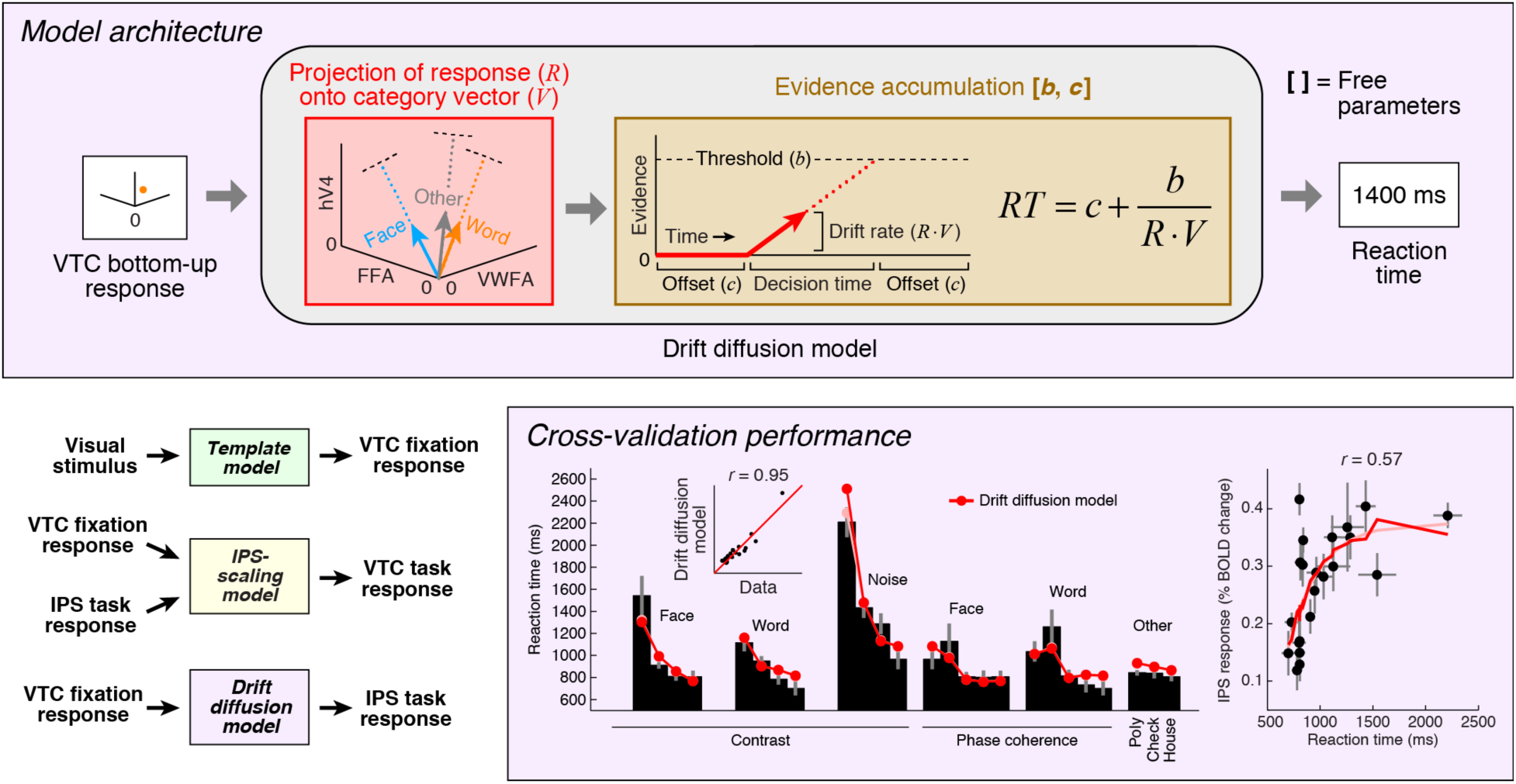
Model of perceptual decision-making in IPS. We implement a model that links the stimulus representation in VTC to a decision-making process occurring in IPS. The model first uses the bottom-up VTC response as a measure of sensory evidence and predicts reaction times in the categorization task (bottom, format same as **Figure 4**). The model then predicts the IPS response as a monotonically increasing function of reaction time (bottom right). For detailed cross-validation results and control models, see **Extended Data Figure 3**. A schematic summarizing all components of our computational model is shown at the bottom left.

Our implementation of the Drift diffusion model involves two steps. First, we use VTC responses during the fixation task (reflecting sensory evidence) to predict reaction times measured in the categorization task. The quality of the predictions is quite high (**Figure 5**, **lower purple box, left**). Second, we apply a simple monotonic function to the reaction times measured during the categorization task to predict the level of response in the IPS (see Methods). The rationale is that the IPS response is sustained over the duration of the decision-making process^46^, and so the overall response integrated over time should be larger for longer decisions. The cross-validated predictions explain substantial variance in IPS (**Figure 5**, **lower purple box, right**).

It is possible to offer a psychological explanation of IPS activity as reflecting cognitive effort— for example, we can posit that IPS activity is enhanced for low-contrast stimuli because the observer works harder to perceive these stimuli. The value of the model we have proposed is that it provides a quantitative and formal explanation of the computations that underlie “effort”. According to the model, categorization of low-contrast stimuli is difficult because the IPS computations required to perform the task involve longer accumulation time. This relates the cognitive task performed by the subject to IPS activity, and suggests that top-down modulation of VTC is a direct consequence of the fulfillment of task demands. We have substantiated this hypothesis for the categorization task and posit that this will serve as a foundation for modeling more complex cognitive tasks such as one-back.

## Discussion

In summary, we have measured and modeled how bottom-up and top-down factors shape responses in VTC. A template operation on low-level visual properties generates a bottom-up stimulus representation, while top-down modulation from the IPS scales this representation in service of the behavioral goals of the observer. We develop a computational approach that posits explicit models of the information processing performed by a network of interacting sensory and cognitive regions of the brain and validate this model on experimental data. The model we have developed stands as the first computational explanation of word-and face-selective responses in VTC, defining a benchmark for future research to build on. We make publicly available data and open-source software code implementing the model at [URL will be inserted upon publication].

The fact that cognitive factors substantially affect stimulus representation in visual cortex highlights the importance of tightly controlling and manipulating cognitive state when investigating stimulus selectivity. In the present measurements, the most striking example comes from stimulus contrast. When subjects perform the fixation task, the contrast-response function (CRF) in VWFA is monotonically increasing, whereas during the one-back task, the CRF flips in sign and is monotonically decreasing (see **Figure 1**). This effect (similar to what is reported in ref. 47) is puzzling if we interpret the CRFs as indicating sensitivity to the stimulus contrast, but is sensible if we interpret the CRFs as instead reflecting the interaction of stimulus properties and cognitive processes. The influence of cognition on visual responses forces us to reconsider studies that report unexpected tuning properties in VTC and IPS, such as tuning to linguistic properties of text in VWFA^48^ and object selectivity in parietal cortex^49^. In experiments that do not tightly control the cognitive processes executed by the observer, it is impossible to distinguish sensory effects from cognitive effects. Our quantitative model of VTC-IPS interactions provides a principled baseline on which to re-interpret past findings, design follow-up experiments, and guide data analysis.

There are a number of research questions that remain unresolved. First, our working hypothesis is that sensory information arrives at VTC, these signals are routed to IPS for evidence accumulation, and then feedback from the IPS modulates the VTC response. Now that this theory is formalized as a series of computations, we can use temporally resolved measurements of neural activity (e.g., EEG, MEG, ECoG) to measure the dynamics of VTC-IPS interactions and extend model predictions into the temporal domain. Second, the computational model presented here is accurate but by no means perfect. For example, the Template model does not capture the step-like response profile of VTC as phase coherence is varied (see **Figure 4**). By exploring a larger range of stimuli and designing experiments to test specific components of the model, we can develop a more nuanced model of stimulus selectivity. Finally, IPS is part of larger brain networks involved in attention^50^ and decision-making^44^, and identifying the computational roles of other regions in these networks is necessary for a comprehensive understanding of the neural mechanisms of perception.

## Acknowledgments

We thank K. Grill-Spector for providing the face and house stimuli used in the main experiment, R. Kiani and N. Kriegeskorte for providing the object stimuli used in the retinotopic mapping experiment, A. Vu and E. Yacoub for collecting pilot data, C. Gratton, M. Harms, and L. Ramsey for scanning assistance, K. Weiner for assistance with ROI definition, and P. Elder for contributions to the development of the model. We also thank C. Gratton, S. Petersen, A. Rokem, A. Vogel, and J. Winawer for helpful discussions. This work was supported by the McDonnell Center for Systems Neuroscience and Arts & Sciences at Washington University (K.N.K.). Computations were performed using the facilities of the Washington University Center for High Performance Computing, which were partially provided through grant NCRR 1S10RR022984-01A1.

## Author Contributions

K.N.K. and J.Y. conceived and designed the experiments. K.N.K. conducted the experiment and analyzed the functional and behavioral data. J.Y. analyzed the diffusion data. K.N.K. and J.Y. wrote the paper.

## Author Information

The authors declare no competing interests.

## METHODS

**<u>Subjects</u>**

Eleven subjects participated in this study. Two subjects were excluded due to inability to identify VWFA in one subject and low signal-to-noise ratio in another subject, leaving a total of nine usable subjects (age range 25–32; six males, three females). All subjects were healthy right-handed monolingual native-English speakers, had normal or corrected-to-normal visual acuity, and were naive to the purposes of the experiment. Informed written consent was obtained from all subjects, and the experimental protocol was approved by the Washington University in St. Louis Institutional Review Board. Each subject participated in 1–3 scanning sessions, over the course of which anatomical data (T1-weighted high-resolution anatomical volume, diffusion-weighted MRI data) and functional data (retinotopic mapping, functional localizer, main experiment) were collected.

### Visual stimuli

Stimuli were presented using an NEC NP-V260X projector. The projected image was focused onto a backprojection screen and subjects viewed this screen via a mirror mounted on the RF coil. The projector operated at a resolution of 1024 × 768 at 60 Hz, and the viewing distance was 340 cm. A Macintosh laptop controlled stimulus presentation using code based on Psychophysics Toolbox^51,52^. Approximate gamma correction was performed by taking the square root of pixel intensity values before stimulus presentation. Behavioral responses were recorded using a button box.

The experiment consisted of 22 types of stimuli. All stimuli were small grayscale images (approximately 2° × 2°) presented at fixation. Each stimulus type consisted of 10 distinct images (e.g. 10 different faces for a face stimulus), and a subset of these images were presented on each given trial.

*FACE (1 stimulus)*.This stimulus consisted of a face pictured from a frontal viewpoint. Ten distinct faces were prepared. Faces were masked using a circle with diameter 2°. The outer 0.25° of the mask was smoothly ramped using a cosine function.

*WORD (1 stimulus)*. This stimulus consisted of a 5-letter word. Ten distinct words were prepared. Letters were white on a gray background, generated using the Helvetica font, and occupied a rectangular region measuring 3.15° × 1.05°.

*PHASE COHERENCE (8 stimuli)*. These stimuli consisted of the FACE and WORD stimuli prepared at four phase-coherence levels: 0%, 25%, 50%, and 75%. To achieve this, for each of the ten images from each stimulus type, the portion of the image within a fixed region (FACE: 2° × 2° square; WORD: 3.15° × 1.05° rectangle) was extracted, and its phase spectrum was blended, to different degrees, with a randomly generated phase spectrum. For example, 25% coherence indicates that the phase of each Fourier component was set to a value that lies at 75% of the angular distance from the original phase to the phase in the randomly generated spectrum.

*NOISE (0 stimuli)*. This stimulus is the FACE stimulus at 0% phase coherence.

*CONTRAST (9 stimuli)*. These stimuli consisted of the FACE, WORD, and NOISE stimuli prepared at three contrast levels. The contrasts of the original stimuli were taken to be 100%, and different contrast levels were achieved by scaling pixel intensity values towards the background value. Contrast levels of 4%, 6%, and 10% were used for the FACE and NOISE stimuli, and contrast levels of 3%, 5%, and 8% were used for the WORD stimulus. This choice of contrast levels matches the average root-mean-square (RMS) contrast across stimulus types (e.g., the average RMS contrast for the FACE stimulus at 4% contrast is approximately equal to the average RMS contrast for the WORD stimulus at 3% contrast). Note that a contrast level of 0% was achieved by estimating responses to blank trials (see *GLM analysis*).

*POLYGON (1 stimulus)*. This stimulus consisted of a string of three polygons (each chosen randomly from a set of polygons). Polygons were white, unfilled, on a gray background, and occupied a region similar in size to that of the WORD stimulus. Ten distinct strings were prepared.

*CHECKERBOARD (1 stimulus)*. This stimulus consisted of alternating black and white square checks. Ten checkerboards were prepared by varying check size from 0.03125° to 0.5° using ten equally spaced steps on a logarithmic scale. The *x*- and *y*-positions of each checkerboard were set randomly. Checkerboards were masked using a circle with diameter 2°.

*HOUSE (1 stimulus)*. This stimulus consisted of a house pictured from a frontal viewpoint. Ten distinct houses were prepared. Houses were masked using a 2° × 2° square. The outer 0.25° of the mask was smoothly ramped using a cosine function.

### Experimental design and tasks

Stimuli were presented in 4-s trials, one stimulus per trial. In a trial, four images from a given stimulus type (e.g. FACE, 10% contrast) were presented sequentially using an 800-ms ON, 200-ms OFF duty cycle. To generate the sequence of four images, we first randomly selected four distinct images out of the ten images associated with the stimulus type. Then, for certain trials (details below), we modified the sequence to include a repetition by randomly selecting one of the images (excluding the first) and replacing that image with the previous image. Throughout stimulus presentation, a small dot (0.12° × 0.12°) was present at the center of the display. The dot switched to a new randomly selected color every 600 ms using a set of six possible colors: magenta, red, yellow, green, cyan, and blue.

In the experiment, two of the stimuli were duplicated (FACE and WORD), yielding a total of 24 stimulus conditions. Data corresponding to these duplicate stimuli are not used in this paper. Each run began and ended with a 16-s baseline period in which no stimuli were presented. During a run, each of the 24 stimulus conditions was presented three times. Six blank trials (no stimulus) were also included. The order of stimulus and blank trials was random, subject to the constraints that blank trials could not occur first nor last, blank trials could not occur consecutively, and no stimulus condition could occur consecutively. During the baseline periods and blank trials, the small central dot was still present. A randomly selected two of the three trials associated with each stimulus condition were modified to include an image repetition (as described previously). Each run lasted 344 seconds (5.7 min).

For each run, subjects were instructed to maintain fixation on the central dot while performing one of three tasks. In the *fixation task*, subjects were instructed to press a button whenever the central dot turned red, and were additionally reminded to not confuse the red and magenta colors. In the *categorization task*, subjects were instructed to report for each stimulus trial whether they perceived a word, a face, or neither (“other”). Responses were made using three different buttons, and subjects were reminded to make only one response for each 4-s trial. In the *one-back task*, subjects were instructed to press a button whenever an image was repeated twice in a row, and were informed that repetitions occurred only within stimulus trials and not across trials. Subjects were warned that although some stimuli are faint (low contrast), they should still try their best to perform the categorization and one-back tasks. Subjects were also informed that some trials are blank trials and that responses were not expected on these trials. Subjects were familiarized with the stimuli and tasks before the actual experiment was conducted.

Subjects performed each of the three tasks four times during the course of the experiment, yielding a total of 3 tasks × 4 runs = 12 runs. The physical stimulus sequence (including the temporal ordering of stimulus images and dot colors) was held constant across tasks. This was accomplished by generating four distinct stimulus sequences and cycling through the sequences and tasks. Specifically, the order of stimulus sequences was ABCD ABCD ABCD, where each letter corresponds to a distinct sequence, and the order of tasks was XYZ XYZ XYZ XYZ, where each letter corresponds to a distinct task. The order of tasks was counterbalanced across subjects. Each stimulus and task combination (e.g. CHECKERBOARD during one-back task) occurred a total of 3 trials × 4 runs = 12 times over the course of the experiment.

### MRI data acquisition

MRI data were collected at the Neuroimaging Laboratory at the Washington University in St. Louis School of Medicine using a modified 3T Siemens Skyra scanner and a 32-channel RF coil. For functional data, 28 oblique slices covering occipitotemporal cortex were defined: slice thickness 2.5 mm, slice gap 0 mm, field-of-view 200 mm × 200 mm, phase-encode direction anterior-posterior. A T2*-weighted, single-shot, gradient-echo EPI sequence was used: matrix size 80 × 80, TR 2 s, TE 30 ms, flip angle 77°, nominal spatial resolution 2.5 mm × 2.5 mm × 2.5 mm. Fieldmaps were acquired for post-hoc correction of EPI spatial distortion. To achieve comprehensive coverage for localization of top-down effects, a whole-brain version of the protocol involving 58 slices and a multiband^53^ factor of 2 was used in three of the nine subjects. In addition to functional data, T1-weighted anatomical data (MPRAGE sequence, 0.8-mm resolution) and diffusion-weighted data (spin-echo EPI sequence, 2-mm resolution, 84 directions, *b*-values of 1,500 and 3,000) were acquired. The diffusion sequence was acquired twice, reversing the phase-encode direction, in order to compensate for spatial distortions. Diffusion data were not acquired for one subject due to time constraints.

### Behavioral analysis

Behavioral results for the categorization task are used in the present study. We analyzed both reaction times (RT) and category judgments. We defined RT as the time elapsed between the onset of the first of the four images in a given trial and the button press. Trials in which no buttons were pressed were ignored. For each subject, we summarized RTs by computing the median RT across trials for each stimulus. To obtain group-averaged RTs, we added a constant to each subject’s RTs in order to match the mean RT to the grand mean across subjects and then computed the mean and standard error across subjects (this normalization procedure compensates for additive offsets in RT across subjects). Category judgments were analyzed by calculating percentages of trials on which a given subject categorized a given stimulus into each of the three categories (word, face, other). Subjects were highly consistent in their judgments: for each stimulus, the most frequently reported category was the same across subjects and was reported more than 85% of the time. Category judgments obtained from the categorization task are used in the labeling and interpretation of experimental results (e.g. **Figures 1**, **2**).

### Diffusion analysis

Subject motion was corrected by co-registering each volume to the average of the non-diffusion-weighted *b* = 0 images. Gradient directions were adjusted to account for the co-registration. From pairs of volumes acquired with reversed phase-encode directions, the susceptibility-induced off-resonance field was estimated using a method similar to that described in ref. 54 as implemented in FSL^55^. Eddy currents were corrected using FSL’s *eddy* tool. The *b* = 3,000 measurements were used to estimate fiber orientation distribution functions for each voxel using constrained spherical deconvolution as implemented in *mrtrix*^56^ (CSD, *l*_*max*_ = 4), and fiber tracts were estimated using probabilistic tractography (500,000 fibers). For each subject, we identified the vertical occipital fasciculus (VOF) using a previously published algorithm^31^, and then quantified the Euclidean distance from the VOF terminations to word-and face-selective regions in VTC and the task-related hotspot in the IPS.

### Pre-processing of anatomical and functional data

The T1-weighted anatomical volume acquired for each subject was processed using FreeSurfer^57^. The results were used to create a cortical surface reconstruction positioned halfway between the pial surface and the boundary between gray and white matter. We used the *fsaverage* surface from FreeSurfer to define anatomical ROIs (details below). These ROIs were transformed to native subject space by performing nearest-neighbor interpolation on the spherical surfaces created by FreeSurfer (these surfaces reflect folding-based alignment of individual subject surfaces to the *fsaverage* surface).

Functional data were pre-processed by performing slice time correction, fieldmap-based spatial undistortion, motion correction, and registration to the subject-native anatomical volume. The combined effects of distortion, motion, and registration were corrected using a single cubic interpolation of the slice time corrected volumes. Interpolations were performed directly at the vertices of the subject’s cortical surface, thereby avoiding unnecessary interpolation and improving spatial resolution^58^.

### GLM analysis

The pre-processed fMRI data were analyzed using GLMdenoise^59^ (http://kendrickkay.net/GLMdenoise/), a data-driven denoising method that derives estimates of correlated noise from the data and incorporates these estimates as nuisance regressors in a general linear model (GLM) analysis of the data. For our experiment, we coded each stimulus and task combination as a separate condition and also included the blank trials, producing a total of (24 stimulus + 1 blank) × 3 tasks = 75 conditions. The response to blank trials was interpreted as the response to a 0%-contrast stimulus. Estimates of BOLD response amplitudes (beta weights) were converted to units of percent BOLD signal change by dividing amplitudes by the mean signal intensity observed at each vertex. To obtain ROI responses, beta weights were averaged across the vertices composing each ROI. Error bars (68% CIs) on beta weights were obtained by bootstrapping runs.

Group-averaged beta weights were calculated by scaling the beta weights obtained for each subject in a given ROI to be a unit-length vector, computing the mean across subjects and bootstrapping subjects to obtain error bars (68% CIs), and rescaling the result such that the mean of the beta weights is equal to the grand mean of the unnormalized beta weights across subjects. This normalization procedure compensates for differences in the overall scale of evoked BOLD responses across subjects and provides interpretable units for the end result. Note that in some cases, beta weights are repeated for easier visualization (e.g., in **Figure 1**, NOISE at 100% contrast is the same data point as FACE at 0% phase coherence).

### Region-of-interest (ROI) definition

Visual field maps were defined using the population receptive field technique applied to retinotopic mapping data^5,37^. Subjects participated in 2–4 runs (300-s each) in which they viewed slowly moving apertures (bars, wedges, rings) filled with a colorful texture of objects, faces, and words placed on an achromatic pink-noise background. The aperture and texture were updated at 5 Hz, and blank periods were included in the design^5^. A semi-transparent fixation grid was superimposed on top of the stimuli^60^. Stimuli occupied a circular region with diameter 10° and the viewing distance was 251 cm. A small semi-transparent central dot (0.15° × 0.15°) was present throughout the experiment and changed color every 1–5 s. Subjects were instructed to maintain fixation on the dot and to press a button whenever its color changed. The time-series data from this experiment were modeled using the Compressive Spatial Summation model^37^ as implemented in analyzePRF (http://kendrickkay.net/analyzePRF/). Angle and eccentricity estimates provided by the model were then visualized on cortical surface reconstructions and used to define V1, V2, V3, and hV4^61^.

Category-selective regions FFA and VWFA were defined using functional localizers^62,63^. Subjects participated in 2 runs (336-s each) in which they viewed blocks of words, faces, abstract objects, and noise patterns. Each block lasted 16 s and consisted of 16 images presented at a rate of 1 Hz. The images differed from those in the main experiment. In each run, the four stimulus types were presented four times each in pseudorandom order, with occasional 16-s blank periods. A semi-transparent fixation grid was superimposed on top of the stimuli^60^. Stimuli occupied a 4° × 4° square region, with the words, faces, and objects occupying the central 3° × 3° of this region. The viewing distance was 340 cm. Subjects were instructed to maintain central fixation and to press a button when the same image is presented twice in a row. The time-series data from this experiment were analyzed using a GLM to estimate the amplitude of the BOLD response to the four stimulus categories.

To define FFA and VWFA, responses to the four stimulus categories were visualized on cortical surface reconstructions. FFA and VWFA were defined based stimulus selectivity, anatomical location, and topological relationship to retinotopic areas^32,63,64^. We defined FFA as face-selective cortex located on the fusiform gyrus. We included in the definition both the posterior fusiform gyrus (pFus-faces/FFA-1) and middle fusiform gyrus (mFus-faces/FFA-2) subdivisions of FFA^64^. We defined VWFA as word-selective cortex located in and around the left occipitotemporal sulcus. In some subjects, multiple word-selective patches were found, and all of these patches were included in the definition of VWFA.

Anatomically-defined ROIs were also created (see **Figures 3a** and **3b**). Based on curvature values on the *fsaverage* surface, we created an anatomical mask of the IPS by selecting the posterior segment of the intraparietal sulcus^65^. Using the atlas of visual topographic organization provided by Wang et *al*.^28^, we estimate that this IPS mask overlaps V3A, V3B, IPS-0, IPS-1, and IPS-2. The locations of IPS-0/1/2/3 from the atlas are shown in **Figure 3** and **Extended Data Figure 2**. We also created an anatomical mask of VTC by computing the union of the *fusiform* and *inferiortemporal* parcels provided by the FreeSurfer Desikan-Killiany atlas^66^ and trimming the anterior extent of the result to include only visually responsive cortex. The VTC mask includes both FFA and VWFA as well as surrounding cortex.

In our data, we find that word-selective visual cortex in some subjects is confined to the left hemisphere, consistent with previous studies^32^. Therefore, to ease interpretation, we restricted our analysis to VWFA, FFA, VTC, and IPS taken from the left hemisphere. In addition, we restricted the definition of V1, V2, V3, hV4, VTC, and IPS to include only vertices exhibiting response amplitudes in the main experiment that are positive on average. This procedure excludes voxels with peripheral receptive fields which typically exhibit negative BOLD responses to centrally presented stimuli.

### Task-based functional connectivity

To identify the cortical region that generates top-down effects in VWFA and FFA, we performed a simple connectivity analysis. First, we averaged BOLD responses across our VTC mask, given that top-down effects appear broadly across VTC. Next, we identified the component of the VTC response that is of no interest, specifically, the bottom-up stimulus-driven response. Our estimate of this component is given by our measurement of VTC responses during the fixation task (22 stimuli + 1 blank = 23 values). We then subtracted the bottom-up component from the VTC response measured during the categorization task (23 values) and one-back task (23 values). This produced a set of residuals (46 values) that reflect the top-down effect in VTC. Finally, we correlated the residuals with the responses of each cortical location in our dataset during the categorization and one-back tasks (46 values). The cortical location that best correlates with the residuals is interpreted as a candidate region that supplies top-down modulation to VTC.

Results were visualized by averaging correlation values across subjects based on the *fsaverage* cortical alignment and plotting results on the *fsaverage* surface. Results from the three subjects for which whole-brain fMRI data were acquired are shown in **Figures 3a** and **3b**. Results from the remaining six subjects with limited fMRI coverage are provided in **Extended Data Figure 2**. Note that the correlation-based analysis we have used is most suitable for connectivity effects that are additive in nature. However, at a finer scale, the nature of the modulation is more accurately characterized as a scaling effect (see **Figure 2**).

There are three important differences between the connectivity analysis described here and conventional correlation-based resting-state functional connectivity (RSFC)^67^ and the psychophysiological interactions (PPI) technique^68^. One is that our connectivity is performed on data that have explicit manipulation of stimulus and task (unlike RSFC). Another is that we analyze the data explicitly in terms of information-processing operations performed by the brain (unlike PPI). In other words, functional connectivity is characterized, not as correlated signal fluctuations, but as a direct consequence of information-processing operations. A third difference is that our connectivity is performed on beta weights that pool across trials^69^, as opposed to raw BOLD time-series. This concentrates the analysis on brain responses that are reliably driven by the stimulus and task, and de-emphasizes trial-to-trial fluctuations in cognitive performance^70^.

### Computational modeling

We developed a computational model to account for BOLD responses measured in VTC and IPS. The model is composed of three components, each of which addresses a different aspect of the data (**Figure 5**, **bottom left**). The first component (*Template model*) specifies how a given stimulus drives bottom-up VTC responses as measured during the fixation task; the second component (*IPS-scaling model*) specifies how top-down modulation from the IPS during the categorization and one-back tasks affects VTC responses; and the third component (*Drift-diffusion model*) specifies how accumulation of evidence from VTC predicts reaction times and IPS responses during the categorization task. Here we describe the modeling approach common to all three components; in later sections we describe each component in greater detail.

Computational modeling was performed on group-averaged beta weights and reaction times using nonlinear least-squares optimization (MATLAB Optimization Toolbox). Leave-one-out cross-validation was used to assess model accuracy (thus, we assess the ability of models to generalize to stimuli that the models have not been trained on). Accuracy was quantified as the percentage of variance explained (*R*^2^) between cross-validated predictions of the data and the actual data. In the case of beta weights, variance was computed relative to 0% BOLD signal change^37^. To assess reliability of cross-validation results, model fitting and cross-validation were repeated for each bootstrap of the group-averaged data (resampling subjects with replacement). For benchmarks on cross-validation performance, we calculated noise ceilings using Monte Carlo simulations^37^ and quantified the performance of a flat-response model that predicts the same response level for each data point.

### Template model

*Basic model description*. The *Template model* specifies the stimulus properties that drive bottom-up responses in VTC. The model accepts as input a grayscale image and produces as output the predicted response in VWFA and FFA during the fixation task. In brief, the model processes the image using a set of V1-like Gabor filters and then computes a normalized dot product between filter outputs and a category template. The category template can be viewed as capturing the prototypical image statistics of a word (VWFA) or face (FFA). The Template model makes no claim as to how the brain might develop category templates; they might be genetically hard-wired^12^ or arise from experience with the environment^71^. The central claim is that the bottom-up information computed by VTC is, at least to first approximation, the output of a template operation applied to the stimulus.

The Template model is related to our previously developed Second-order contrast (SOC) model^38^. Similar to the SOC model, the Template model has a cascade architecture involving two stages of filtering, rectification, and normalization. However, the Template model incorporates a specific second-stage filter (the template), whereas the SOC model uses a variance-like operation in the second stage that captures generic sensitivity to second-order contrast. Whether the Template model captures certain response properties, such as invariance to font in VWFA^11^ or coarse luminance-contrast selectivity in FFA^72^, is an empirical question that can only be resolved through quantitative evaluation on experimental data. For example, our measurements indicate that VWFA responds strongly to polygons (**Figure 1d**); the Template model already accounts for this effect (**Figure 4**, **lower green box**).

*Model details*. The first stage of the Template model involves computing a V1-like representation of the image. The image is first resized to 250 pixels × 250 pixels, and luminance values are mapped to the range [–0.5,0.5], which has the effect of mapping the gray background to 0. The model then calculates V1 energy in the same way as the SOC model^38^. Specifically, the image is projected onto a set of isotropic Gabor filters occurring at 8 orientations, 2 quadrature phases, and a range of positions (63 *x*-positions × 63 *y*-positions). Filters are constructed at a single scale with a peak spatial frequency tuning of 4 cycles per degree (see **Extended Data Figure 3**) and a spatial frequency bandwidth of 1 octave (full-width at half-maximum of the amplitude spectrum). Filters are scaled such that filter responses to full-contrast optimal sinusoidal gratings are equal to one. Outputs of quadrature-phase filters are squared, summed, and square-rooted, analogous to the complex-cell energy model^73^.

After computing V1 energy, the model applies divisive normalization^74^, again analogous to the SOC model. The output of each filter is divided by the average output across filter orientations at the same position:

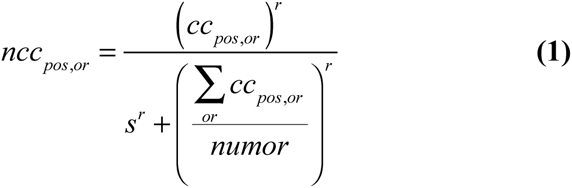

where *ncc*_*pos,or*_ is the normalized filter output at a given position and orientation, *cc*_*pos,or*_ is the filter output at a given position and orientation, *numor* is the total number of orientations, and *r* and *s* are parameters that control the strength of the normalization. For simplicity and to reduce the potential for overfitting, we do not fit *r* and *s* but simply use *r* = 1 and *s* = 0.5, values determined from our previous study^38^.

At this point in the model, the representation of the image is a 3D matrix of dimensions 63 *x*-positions ×63 *y*-positions × 8 orientations. To visualize this representation, a hue-saturation-value image is used (see **Figure 4**, **upper green box**). For each position, a set of 8 vectors is constructed with vector angles corresponding to filter orientation and vector lengths corresponding to normalized filter output. These vectors are averaged and an image pixel is used to summarize the result. Specifically, the hue of a pixel indicates the angle of the vector average and the value of the pixel indicates the length of the vector average.

The second stage of the Template model involves taking the V1-like representation of the image and comparing it to a category template to generate the predicted response. Specifically, the response is computed as

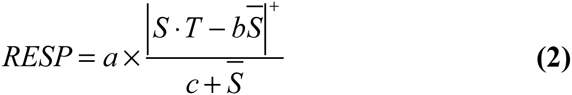

where *RESP* is the predicted response, *S* is the 3D matrix with the V1-like representation of the image, *T* is the category template, 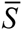 is the average of the elements in *S*, | |^+^ indicates positive half-wave rectification, and *a*, *b*, and *c* are free parameters.

There are three basic steps in **Equation 2**. The first step is a filtering operation, accomplished by computing the dot product between the stimulus and the template (*S·T*). Intuitively, this operation quantifies the similarity between the stimulus and the template. The second step is subtraction of average stimulus energy (– *b* 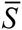) with a free parameter controlling the strength of the subtractive normalization. This subtraction can be interpreted as penalizing non-specific energy in the stimulus, thereby inducing preference for stimulus energy that conforms to the category template. (An alternative interpretation is that the subtraction provides flexibility with respect to the overall mean of the template.) To ease interpretation and ensure that negative responses are not obtained, the result of the subtraction is positively rectified (| |^+^). The third step is division by average stimulus energy (/ (*c*+ 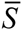)) with a free parameter controlling the strength of the divisive normalization. This division penalizes non-specific energy in the stimulus, similar to subtractive normalization, but induces a different response geometry^75^. In summary, **Equation 2** computes a dot product between the stimulus and the template that is normalized subtractively and divisively by the average stimulus energy.

Where does the category template in **Equation 2** come from? Given that we do not have sufficient sampling of stimuli to directly estimate templates from the data, we adopted the simple strategy of constructing templates from our stimulus set. Specifically, we took the WORD and FACE stimuli at 100% contrast and used the first stage of the Template model to compute a V1-like representation of these stimuli. This produced for each category, ten points in a 63 × 63 × 8 = 31,752-dimensional space. We then computed the centroid of the ten points, producing a category template (example shown in **Figure 4**, **upper green box**). Because the category template is constructed from the same stimuli used in our experiment, it is guaranteed that the Template model predicts large responses to the preferred category (e.g., using a category template constructed from the face stimuli guarantees that the face stimuli produce large responses from the model). However, there is no guarantee that the model will accurately account for responses to the other stimuli used in our experiment.

*Model fitting*. The Template model was fit to the fixation responses of VWFA and FFA. Model outputs were calculated for all ten images associated with a given stimulus type and then averaged to obtain the final model output for that stimulus type. To aid model fitting, the *S·T* and 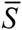 quantities were pre-computed and pre-conditioned by dividing each quantity by the mean of that quantity across stimuli. After pre-conditioning, a variety of initial seeds for *b* and *c* were evaluated in order to avoid local minima. Specifically, we performed optimization starting from initial seeds corresponding to every combination of *b* and *c*, where *b* is chosen from {0.5 1 1.5 2 3 5} and *c* is chosen from {.01.05.1.5 1 5 10}.

*Alternative models*. (1) The *Category model* predicts a fixed response level for stimuli from the preferred stimulus category (word for VWFA, face for FFA) and a different response level for all other stimuli. Category judgments provided by the subjects were used to determine category membership. (2–3) We evaluated simplified versions of the second-stage normalization used in the Template model. One version, *Template model (only subtractive normalization)*, omits the divisive normalization (and thus characterizes responses as a simple linear function of V1-like normalized filter outputs), whereas the other version, *Template model (only divisive normalization)*, omits the subtractive normalization. (3) In *Template model (omit first stage)*, the first stage of the model is omitted and the template operation is performed on a pixel representation of the image (i.e. *S* refers to the original image instead of the V1-like representation of the image). (4–6) We evaluated the effect of using different templates in the Template model. *Template model (non-selective template)* uses a template consisting of all ones. *Template model (mixed template)* uses a template generated by unit-length normalizing both the word and face templates and then averaging the templates together. *Template model (random template)* uses a template generated by drawing uniform random values from the range [0,1].

### IPS-scaling model

*Basic model description*. The *IPS-scaling model* predicts top-down modulation of VTC by taking into account measurements of IPS activity. The model accepts as input the response in VTC (either VWFA or FFA) during the fixation task and the response in IPS during the stimulus-directed tasks (categorization, one-back), and produces as output the predicted response in VTC during the stimulus-directed tasks. Intuitively, the model answers the question: how much is the bottom-up response in VTC enhanced by the IPS when the subject performs a task on the stimulus?

*Model details*. The IPS-scaling model multiplies the bottom-up response in VTC measured during the fixation task by a scaled version of the IPS response observed during a stimulus-directed task:

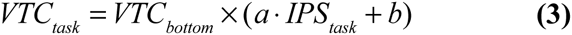

where *VTC*_*task*_ is the predicted response in VTC during the stimulus-directed task, *VTC*_*bottom*_ is the bottom-up response in VTC, *IPS*_*task*_ is the response in IPS during the stimulus-directed task, and *a* and *b* are parameters that allow a scale and offset to be applied to the IPS response. The final scaling factor that is applied to *VTC*_*bottom*_ is shown in **Figure 4**, **bottom yellow box**.

The measurements of IPS activity used in the IPS-scaling model are extracted using a broad anatomical mask of the IPS (see *Region-of-interest (ROI) definition*). This strategy avoids the overfitting that might ensue from a more specific voxel-selection procedure^76^. For example, if we were to select the single cortical location in the IPS that best correlates with the top-down modulation of VTC (see **Figure 3b**), this would make voxel selection a critical part of the model and render the modeling analysis circular.

*Model fitting*. The IPS-scaling model was fit to the fixation, categorization, and one-back responses observed in VWFA and FFA. Leave-one-out cross-validation was performed by systematically leaving out each of the categorization and one-back responses. Since measurement noise is present in the fixation responses, treating the fixation responses as exact estimates of bottom-up responses would result in suboptimal model performance (especially in the case of bottom-up responses that are near zero). We therefore devised a procedure that allows flexibility in estimating bottom-up responses (see light lines in **Figure 4**, **bottom yellow box**). In the procedure, a separate parameter is used to model the bottom-up response associated with each stimulus. During model fitting, these bottom-up parameters are initially set to be equal to the measured fixation responses, parameters of the model excluding the bottom-up parameters are optimized, and then all parameters are optimized simultaneously. This procedure was also used for the alternative models described below.

*Alternative models*. (1) The *Task-invariant model* posits that top-down modulation does not occur and that a fixed set of responses can characterize all three tasks. (2–5) We evaluated several phenomenological models for purposes of comparison. The *Additive model* predicts responses during stimulus-directed tasks by adding a constant to bottom-up responses. The *Scaling model* predicts responses during stimulus-directed tasks by multiplying bottom-up responses by a constant. The *Additive model (task-specific)* and *Scaling model (task-specific)* are identical to the previous two models, except that separate constants are used for the categorization and one-back tasks. (6) The *IPS-additive model* predicts responses during stimulus-directed tasks by adding a scaled version of the IPS response to bottom-up responses in VTC. (7–8) To assess the specificity of the IPS enhancement, we evaluated variants of the IPS-scaling model. In the *IPS-scaling (shuffle)* model, IPS responses are shuffled across stimuli and tasks (restricted to the stimulus-directed tasks) before being used in the model. In the *IPS-scaling (shuffle within task) model*, IPS responses are shuffled across stimuli but not across tasks before being used in the model.

### Drift diffusion model

*Basic model description*. The *Drift diffusion model* specifies the decision-making operations that underlie performance of the categorization task, and is based upon past research on perceptual decision-making^44–46^. The model accepts as input fixation responses in VTC and produces as output predicted reaction times and IPS responses for the categorization task. The basic idea is that VTC responses provide evidence regarding which stimulus category has been presented to the subject, and this evidence is accumulated over time by the IPS in order to make a final decision regarding stimulus category.

*Model details*. First, we collect fixation responses in hV4, VWFA, and FFA and divide each set of responses by their mean. This normalization ensures that different ROIs have similar units. Then, for each stimulus category (word, face, other), we compute the centroid of the fixation responses associated with that category, interpret this centroid as a vector, and normalize the vector to unit length. This procedure generates category vectors, defined in a three-dimensional neural space, that point in the directions of the "arms" of the manifold of the VTC representation (see **Figure 2e**).

Next, we take the VTC fixation response for a given stimulus and project this response onto the category vector associated with that stimulus. The working hypothesis is that this operation is performed by neurons in IPS and that the magnitude of the projection indicates the strength of evidence for that specific category. For example, there might be an IPS neuron that receives information from VTC and responds strongly when the VTC response is consistent with the category vector corresponding to a word.

In accordance with drift diffusion models, we posit that evidence is accumulated until a threshold is reached, at which point the decision is made. This generates a prediction of the reaction time required to perform the categorization task on the stimulus:

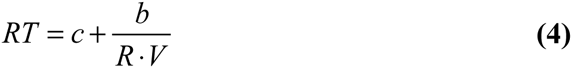

where *RT* is the predicted reaction time, *R* is the VTC fixation response, *V* is the category vector associated with the stimulus, *b* is a parameter that controls the threshold, and *c* is a parameter that compensates for non-decision time (e.g. motor response). *R.V* is interpreted as a drift rate, and *b*/(*R.V*) is the time required to reach the threshold (see **Figure 5**, **upper purple box**).

Given that neuronal responses in parietal cortex reflect the duration of the decision-making process^46^, we can use RT to predict IPS activity. A detailed model relating RT to BOLD measurements of IPS activity requires precise characterization of neural dynamics during decision-making and IPS subdivisions that might represent evidence accumulation for different stimulus categories. For the purposes of this study, we use a simple model that posits a monotonically increasing relationship between RT and the IPS response:

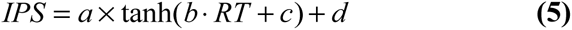

where *IPS* is the predicted IPS response, *RT* is the observed reaction time for a given stimulus, tanh is the hyperbolic tangent function intended as a generic sigmoidal nonlinearity, and *a*, *b*, *c*, and *d* are free parameters.

*Alternative models*. (1) The *Drift diffusion model (separate thresholds)* uses a separate threshold parameter for each stimulus category. This allows us to assess the validity of having a single threshold parameter in the model. (2) The *Drift diffusion model (axis-aligned category vectors)* uses category vectors that are aligned with the axes of the multi-dimensional neural space. For example, in this model, the word category vector is a vector that is one along the VWFA axis and zero along the hV4 and FFA axes. This model tests the idea that evidence for words and faces is contributed only by the VTC regions selective for those categories.

### Code availability

Software code implementing the model proposed in this paper is available at [URL will be inserted upon publication].

---

**Extended Data Figure 1.**
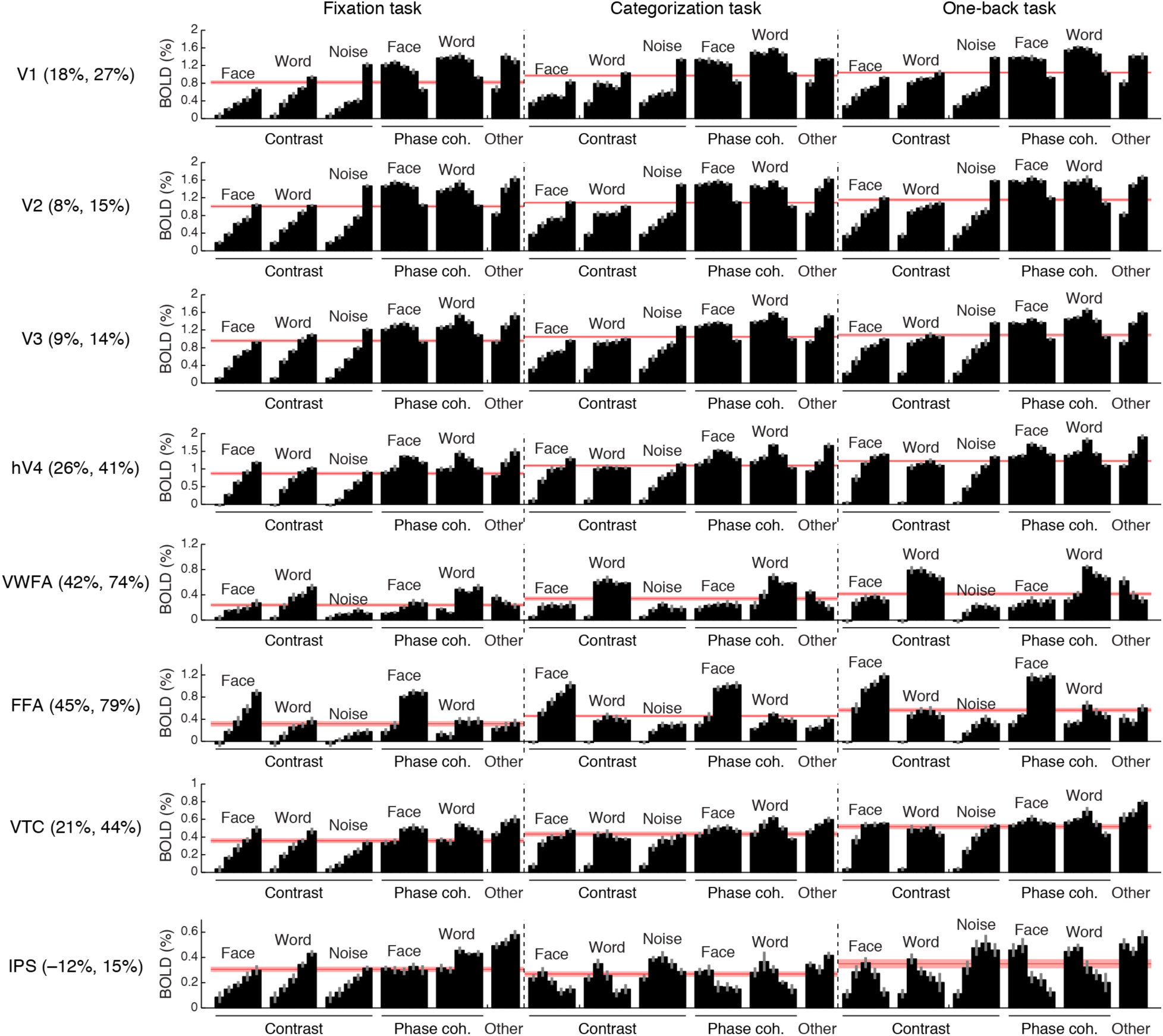
Comprehensive summary of fMRI measurements. Black bars indicate responses (beta weights) evoked by different stimuli and tasks. Red lines indicate the average response across stimuli, computed separately for each task. Error bars indicate bootstrapped 68% CIs (resampling subjects with replacement). Percentages in ROI labels indicate the strength of the response observed during the categorization and one-back tasks relative to the fixation task. For example, in FFA, the average response across stimuli during the one-back task is 79% stronger than the average response across stimuli during the fixation task. Task effects are substantially stronger in VWFA and FFA than in early visual areas V1–V3. The larger apparent task modulation in V1 compared to V2 and V3 might due to small eye movements that may have been made during the categorization and one-back tasks. Our interpretation of the observed IPS activity during the fixation task is that this activity reflects the decision-making process involved in judging the color of the fixation dot. Support for this interpretation comes from the fact that the root-mean-square contrast of the stimuli, computed over a small region surrounding the fixation dot (0.36° × 0.36°), correlates strongly with IPS responses during the fixation task (*r* = 0.86).

**Extended Data Figure 2.**
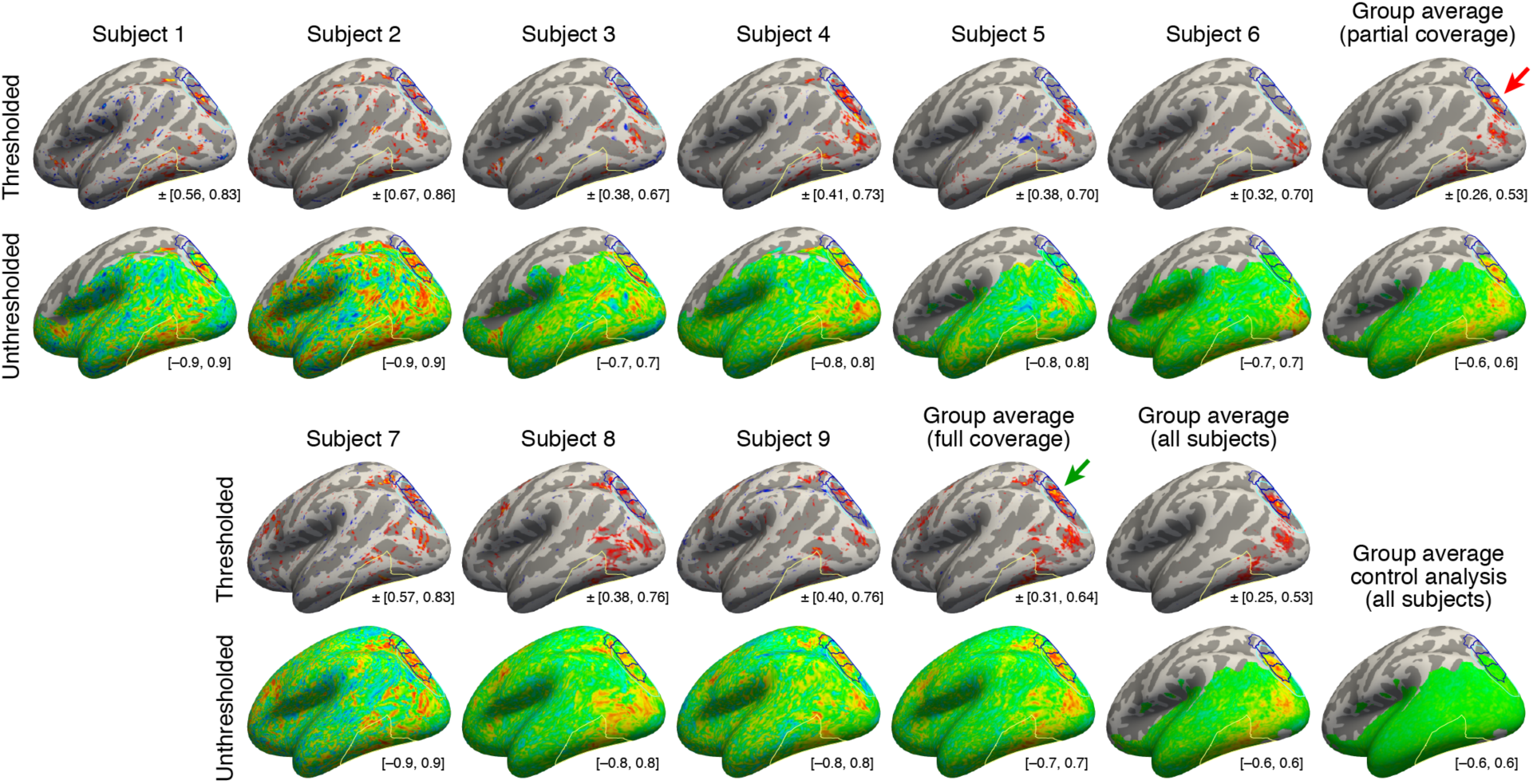
Maps of top-down connectivity to VTC. This figure shows thresholded and unthresholded maps for individual subjects and group averages (same format as **Figure 3b**; all maps shown on the *fsaverage* surface). At the lower right of each map is the range of values used for the colormap. The upper two rows show results obtained for the six subjects with partial brain coverage. Group average results for these subjects are shown in the last column. The lower two rows show results obtained for the three subjects with full brain coverage. Group average results for these subjects are shown in the third to last column. Group average results for all subjects are shown in the second to last column. The bottom right shows results obtained from a control analysis in which we generate individual-subject maps by correlating cortical responses with random Gaussian noise and then average these maps across subjects. This control analysis produces no substantial correlations. Notice that the peak correlation is found in and around IPS-0/1 for both the group of subjects with partial brain coverage (red arrow) and the group of subjects with full brain coverage (green arrow). Some variability in the location of the peak correlation is expected given that there are limits on the degree to which functional areas can be aligned across subjects based solely on anatomical features.

**Extended Data Figure 3.**
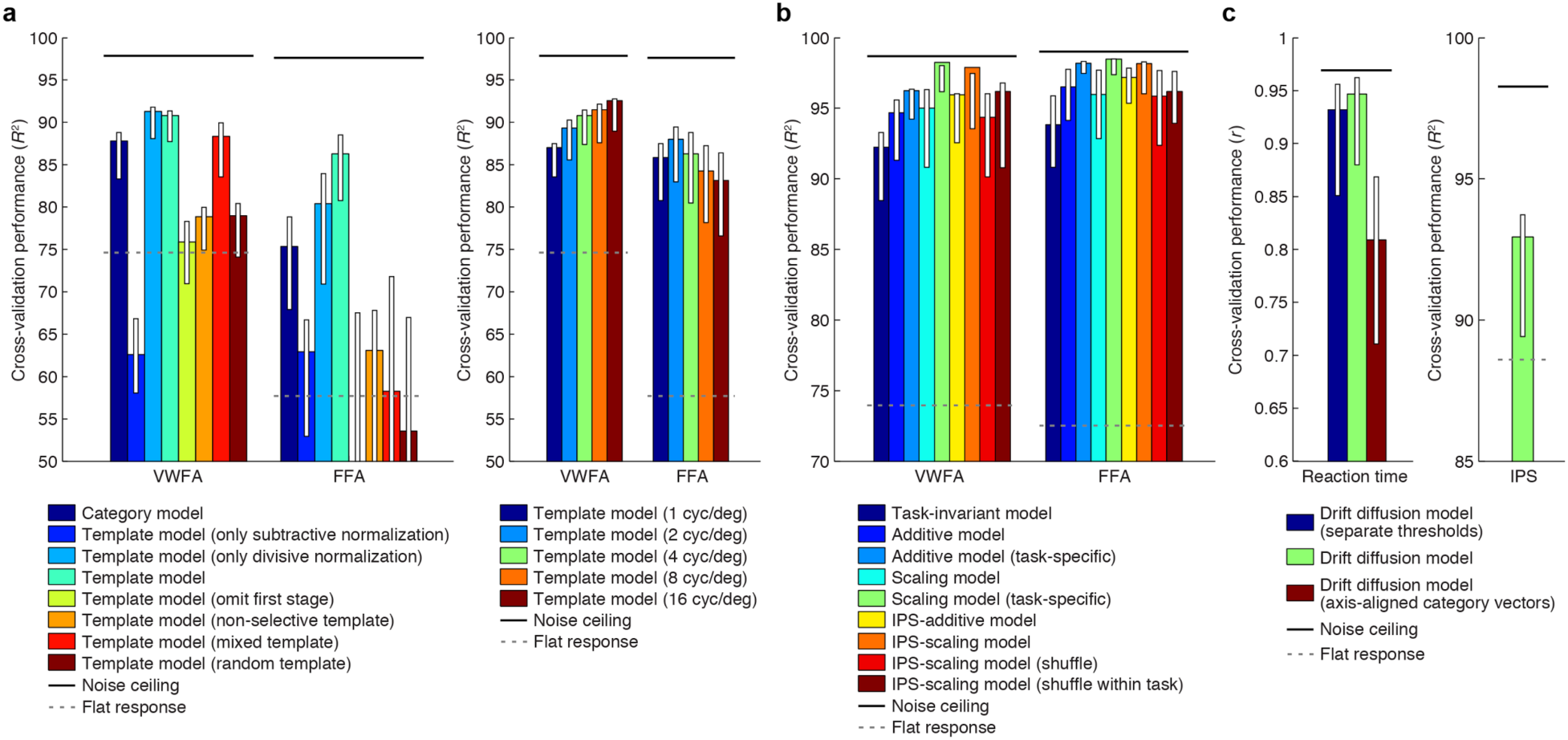
Comparison of performance against control models. Bars indicate leave-one-out cross-validation performance. Error bars indicate 68% CIs, obtained by bootstrapping (resampling subjects with replacement). Solid horizontal lines indicate the noise ceiling, i.e., the maximum possible performance given measurement variability in the data. Dotted horizontal lines indicate the cross-validation performance of a model that predicts the same response level for each data point (this corresponds to *R*^2^ = 0 in the conventional definition of *R*^2^ where variance is computed relative to the mean). (a) *Bottom-up stimulus-driven responses in VTC* (see Figure 4, green panels). The plot to the left shows that the performance of the Template model degrades if the second stage of nonlinearities is omitted (Template model (only subtractive normalization)) or if the first stage of the model involving V1-like filtering is omitted (Template model (omit first stage)). The plot also shows that the precise configuration of the template is important for achieving high model performance (Template model (non-selective, mixed, random templates)). The plot to the right shows performance of the Template model as a function of the spatial frequency tuning of the filters (spatial frequency bandwidth fixed at 1 octave). For simplicity, we use in the Template model a single set of filters that have a spatial frequency tuning of 4 cycles/degree. (b) *Top-down modulation of VTC responses* (see Figure 4, yellow panels). Performance of the IPS-scaling model degrades if the IPS input into the model is shuffled across conditions (IPS-scaling model (shuffle, shuffle within task)), indicating that top-down modulation from the IPS is dependent on the stimulus and task. Although the Additive and Scaling models perform well, note that these are *ad hoc*, phenomenological models. For instance, the Scaling model (task-specific) posits separate parameters for the amount of scaling under the categorization and one-back tasks. However, such a model does not explain *why* there is a different amount of scaling, whereas the IPS-scaling model provides such an explanation. (c) *Reaction times (left) and IPS responses (right) during categorization task* (see Figure 5, purple panels). Performance of the Drift diffusion model does not degrade substantially if a single threshold is used, thus justifying this simplification. Performance degrades if axis-aligned category vectors are used, supporting the assertion that responses of multiple VTC regions are used by subjects in deciding image category.

